# Genome-wide profiling of genetic variation at tandem repeat from long reads

**DOI:** 10.1101/2024.01.20.576266

**Authors:** Helyaneh Ziaei Jam, Justin M. Zook, Sara Javadzadeh, Jonghun Park, Aarushi Sehgal, Melissa Gymrek

## Abstract

Tandem repeats are frequent across the human genome, and variation in repeat length has been linked to a variety of traits. Recent improvements in long read sequencing technologies have the potential to greatly improve TR analysis, especially for long or complex repeats. Here we introduce LongTR, which accurately genotypes tandem repeats from high fidelity long reads available from both PacBio and Oxford Nanopore Technologies. LongTR is freely available at https://github.com/gymrek-lab/longtr.

## Main

Tandem repeats (TRs), including short tandem repeats (STRs; repeat unit 1-6bp) and variable number tandem repeats (VNTRs; repeat unit 7+bp), refer to regions of the genome that consist of adjacent repeated units. TRs are a large source of genetic variation in humans^1^ and are implicated in a growing list of Mendelian and complex traits^2^. In the last decade, multiple tools have been developed to estimate the repeat length and/or sequence of TRs using short reads (e.g.^3–6^) but certain repeats such as highly complex TRs have remained intractable. Long-read sequencing technologies offer a promising solution. However, tools designed for short reads are ineffective on long reads given the considerable differences including in read length, base calling accuracy, error profiles at STRs (**Supplementary Fig. 1**) and paired-end vs. single-end format.

Here, we introduce LongTR, which extends the HipSTR^3^ method originally developed for short read STR analysis in order to genotype STRs and VNTRs from accurate long reads available for both PacBio^7^ and Oxford Nanopore Technologies^8^ (ONT). LongTR takes as input sequence alignments for one or more samples and a reference set of TRs and outputs the inferred sequence and length of each allele at each locus. It uses a clustering strategy combined with partial order alignment to infer consensus haplotypes from error-prone reads, followed by sequence realignment using a Hidden Markov Model, which is used to score each possible diploid genotype at each locus (**Methods; Supplementary Fig. 2**). Unlike other existing long read TR genotypers, LongTR supports multi-sample calling, incorporates read phase information when available, employs a technology-specific homopolymer error model, and outputs genotype quality scores.

We focused on benchmarking LongTR against TRGT^9^, a recently developed TR genotyper that outperforms previous methods^10,11^ on PacBio HiFi reads. We used both tools to genotype TRs from 30x HiFi reads for the well-characterized sample HG002 using the TRGT reference set of 937,122 human TRs. LongTR was 14% faster, taking 569 minutes to complete the genotyping on a single thread, compared to 662 minutes for TRGT. TRGT and LongTR genotyped 99.83% and 99.78% of the repeats respectively. At sites called by both methods, 86.4% of alleles have identical lengths and 98% differ in length by at most a single copy number.

To evaluate the accuracy of genotype calls, we extracted alleles for genotyped TRs from the haplotype-resolved genome assembly of HG002^12^, which was generated using multiple technologies and orthogonal computational methods, and is thus treated as a ground truth here. Of 814,319 repeats for which exactly two alleles were extracted from the assembly, TRGT and LongTR showed 97.6% and 98.3% length concordance allowing for 1 bp differences. LongTR showed similar gains when evaluating sequence concordance. In both cases, the advantage of LongTR over TRGT was highest at long (>500bp) repeats (**Fig. 1**). We identified multiple scenarios in which LongTR calls match the assembly and TRGT does not, including regions with a high number of truncated reads or regions called as large insertions by TRGT that had low read support. Further, LongTR detected 514 TRs with large structural deletions that remove the entire repeat, resulting in a null allele, whereas these cases are not reported by TRGT (**Supplementary Fig. 3**). Overall, these results suggest both tools perform similarly at TRs in regions that are easier to genotype, whereas LongTR obtains higher quality genotypes at longer or structurally complex regions. Notably, we observed multiple instances where both LongTR and TRGT reported identical genotypes that differed from the assembly. These cases usually occurred in complex genomic regions such as large insertions or segmental duplications or regions of high homozygosity, which are known to pose challenges to diploid assemblies^13^ and thus likely represent assembly errors rather than errors from LongTR or TRGT (**Supplementary Fig. 4**).

**Figure 1:**
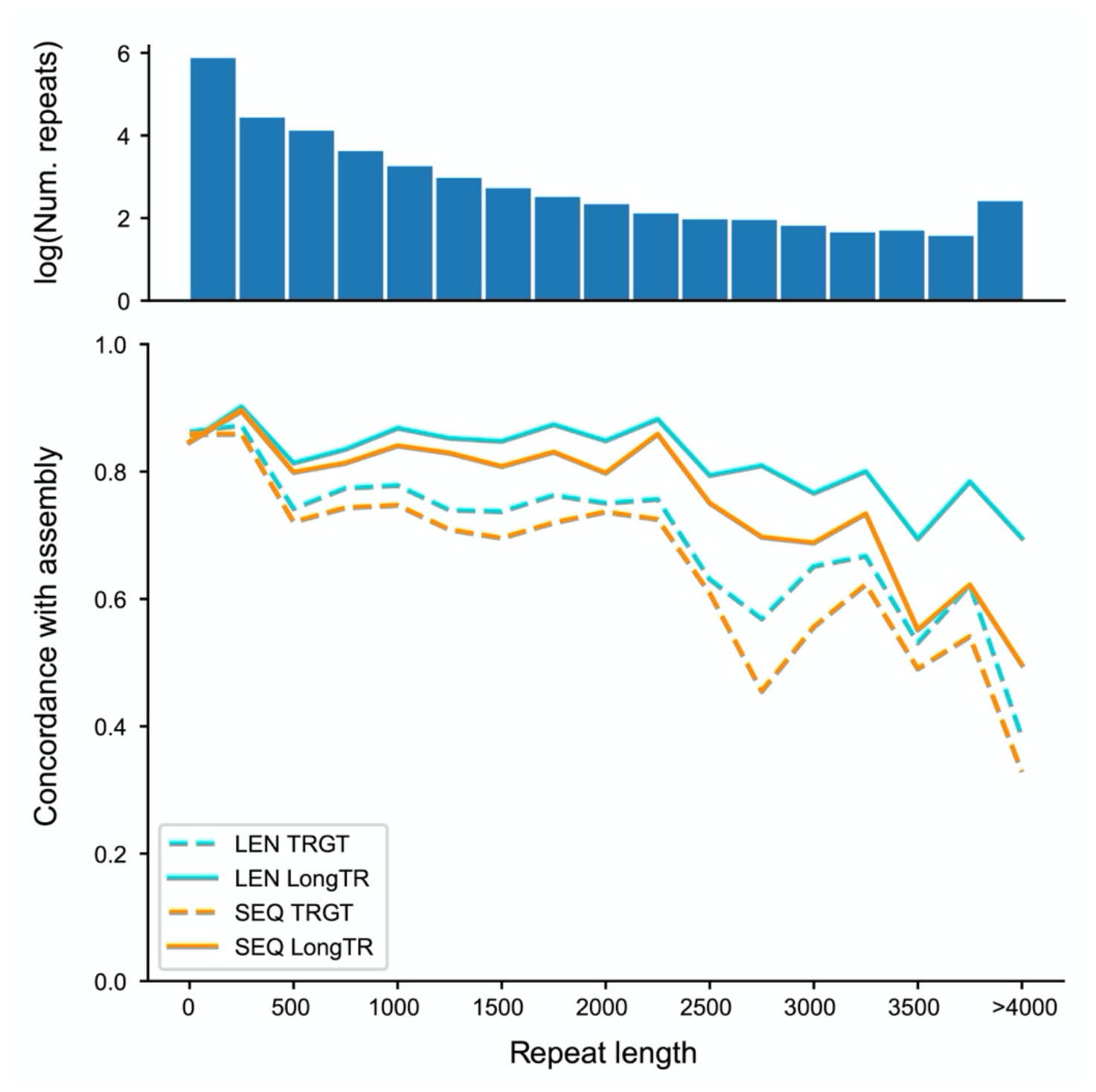
Concordance of TR genotypes obtained from PacBio HiFi with assembly alleles in HG002. TRs were binned by length of the repeat (in bp, bin size = 250bp) in GRCh38. The x-axis shows the TR length and the y-axis shows the percent of alleles that match the assembly. Blue lines show when only length is considered. Orange lines show when both length and sequence are considered. dashed=TRGT; solid=LongTR. The top panel shows the number of repeats in each bin, on a logarithmic scale.

To further evaluate the accuracy of LongTR, we performed TR genotyping in an Ashkenazi trio (HG002, HG003, and HG004) and determined the Mendelian inheritance (MI) of TR genotypes. Overall, LongTR showed 87% MI (compared to 79% for TRGT) at sites where at least one trio member was not homozygous for the reference allele. Mendelian consistency monotonically increases with genotype quality scores computed by LongTR, which can be used to filter low quality calls (**Fig. 2**). Homopolymer repeats showed the lowest consistency with MI of 79.5% for LongTR and 73.9% for TRGT. We trained a PacBio HiFi homopolymer error model at 840,248 homopolymer repeats from the HipSTR reference, which contains homopolymers with more precise boundaries than the TRGT set, by comparing observed repeat lengths at HiFi reads to those obtained from the HG002 assembly (**Supplementary Table 1**). In the Ashkenazi trio we observed an MI of 83% and 81% for LongTR with and without error modeling, and 78% for TRGT at these loci (**Supplementary Fig. 5**). Further, we observed a 13% increase in concordance of LongTR alleles with the HG002 assembly when using the homopolymer error model.

**Figure 2:**
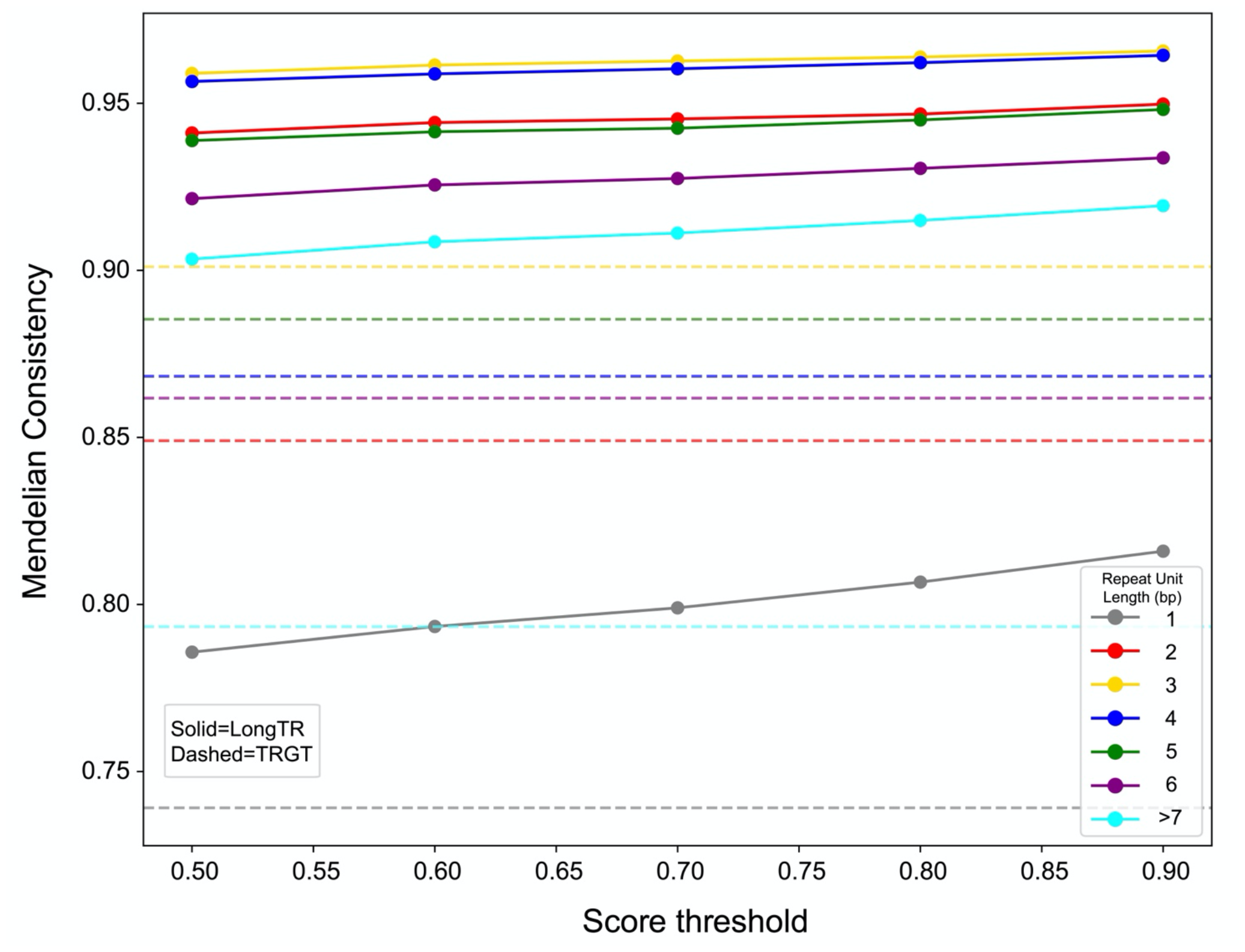
Assessing Mendelian consistency of TR calls in an Ashkenazi trio using PacBio HiFi reads. The x-axis gives the LongTR score threshold to include calls and the y-axis gives the percentage of TRs for which genotypes in the trio follow Mendelian consistency. Trio-TR pairs for which all members were called as homozygous for the reference allele were excluded. Dashed=TRGT; solid=LongTR. Note TRGT does not report a quality score and thus a single horizontal line is shown. Color indicates the size of the repeat unit (in bp) considered.

We sought to further assess the ability of LongTR to genotype longer repeats. First, we compared LongTR genotypes in HG002 to those from adVNTR^14^, a tool specifically designed for genotyping VNTRs, on a reference set of 10,186 autosomal gene-proximal VNTRs. When allowing for differences in up to a single copy number (**Methods**), LongTR and adVNTR showed 96% concordance. adVNTR required approximately 23 hours to genotype HG002, compared to 1.5 hours for LongTR on this repeat set. Second, we determined whether LongTR could identify large expansions in HiFi reads obtained from patients harboring long pathogenic alleles implicated in Huntington’s disease (n=4) and Fragile X Syndrome (n=3). LongTR correctly identified expansions in *HTT* and *FMR1*, including alleles consisting of up to several thousand bp for both loci (**Supplementary Table 2**). Allele sequences reported by LongTR match repeat unit copy number counts for these reads reported on the dataset website for the tested samples (**Data Availability, Supplementary Fig. 6**).

We next evaluated the ability of LongTR to genotype TRs in a separate long read technology, using ONT’s recently released Duplex reads (average length 27kb compared to 15-20kb for PacBio HiFi reads) available for HG002. Overall, we observed high concordance between genotypes obtained from ONT duplex and PacBio HiFi (88% allowing for 1bp off; **Supplementary Fig. 7**) with the latter showing higher concordance with the assembly (99% for PacBio HiFi vs 87% for ONT, allowing for 1bp off). However, we identified several large repeat expansions uniquely detected by the ONT data, which were consistent with the alleles in the assembly. Examining these expansions showed that discrepancies often occurred in regions where few or no PacBio HiFi reads aligned to or spanned the insertion, leading LongTR to inaccurately genotype the locus (**Supplementary Fig. 8**). Our observations suggest that the longer read lengths of ONT enhance the detection accuracy particularly for regions with large insertions compared to the reference.

Finally, we compared TR genotypes obtained from HipSTR on Illumina short reads and LongTR on PacBio HiFi reads for HG002. For this analysis, we used the hg38 reference set of 1,638,945 STRs available from HipSTR’s website, of which only 1.6% are longer than 100bp. Of 1,556,278 STRs that were genotyped by both methods, 88% were concordant, increasing to 97% if allowing for 1bp length difference (**Supplementary Fig. 9a**). HipSTR reported homozygous reference for all repeats with length above 250bp (the Illumina read length) and no read distribution information, indicating genotypes for longer repeats are not reliable. For shorter repeats, concordance (by allowing 1bp off) decreases with increasing length of the repeat (**Supplementary Fig. 9b**).

Overall, LongTR provides accurate sequence-resolved TR genotyping from long reads for nearly 99% of TRs in mappable regions of the genome and outperforms existing methods for this task. While the majority of TRs can be resolved, we identified multiple TRs with large, complex insertions relative to the reference that are challenging to span even with long reads and may in some cases be misrepresented by reference assemblies. These cases may represent the limits of mapping-based approaches. Future work is needed to incorporate alternative approaches, such as pangenome or assembly-based methods, that do not suffer from these limitations. We envision these improvements will enable systematic incorporation of TRs into genome-wide analyses for a range of applications.

## Acknowledgements

This work was supported in part by NIH grants 1R01HG010149 (M.G.) and U01DA051234 (M.G.). Certain commercial equipment, instruments, or materials are identified to specify adequately experimental conditions or reported results. Such identification does not imply recommendation or endorsement by the National Institute of Standards and Technology, nor does it imply that the equipment, instruments, or materials identified are necessarily the best available for the purpose. We thank Eric Mendenhall for helpful comments on the manuscript.

## Author contributions

H.Z.J. developed LongTR, led analyses, and wrote the manuscript. J.M.Z. helped perform comparison of LongTR and TRGT genotypes to the HG002 assembly and helped supervise ONT analyses. S.J. and J.P. helped with adVNTR analyses. A.S. helped with testing and development of LongTR. M.G. supervised development of LongTR and analyses and wrote the manuscript.

## Methods

### Overview of the LongTR method

LongTR extends HipSTR^3^, which was originally developed to analyze short reads, to genotype both VNTRs and STRs using accurate long reads. Here, we use accurate long reads to refer to PacBio HiFi and ONT duplex reads, each of which have been shown to have per-base error rates comparable to those of Illumina reads^15,16^. Like HipSTR, LongTR begins with aligned reads for one or more samples and the reference coordinates for a predefined set of TRs. Then for each TR, it extracts reads encompassing the repeat, infers candidate TR haplotypes, and uses a Hidden Markov Model to realign all the reads overlapping a repeat region to the candidate haplotypes. Finally, it outputs a VCF file containing each individual’s TR genotypes and corresponding quality scores (**Supplementary Fig. 2**). Below we discuss key steps where LongTR differs from the HipSTR method to enable analysis of TRs from long reads.

#### Haplotype identification

LongTR uses a new method for the identification of candidate haplotypes from input reads of samples at each TR. It starts by trimming all reads aligned to the target TR of interest to include the repeat plus a user-defined window of context sequence. It then iterates over all trimmed sequences and includes any sequence supported by a sufficient number of reads by one or more samples (at least two reads and more than 20% of reads in a single sample, or more than 5% of all reads across all samples) as a candidate haplotype.

In some scenarios, typically when the repeat is long or the region is complex with multiple insertions and deletions, reads from a haplotype fail to meet the criteria set in the first stage, resulting in their exclusion from the set of haplotypes. To address this, we perform a second iteration during which we identify samples for which over 25% of aligned reads lack a corresponding representative haplotype and form additional candidate haplotypes using excluded reads for each of these samples. These previously excluded reads for each sample are then sorted by length of the sequence aligned to the repeat region, after which a greedy clustering algorithm is applied to form the initial sequence clusters. In this method, the first cluster is formed by designating the first sequence as its centroid. Starting with the second sequence *S* we evaluate whether there exists a centroid *C* within the centroids set for which the edit distance between *S* and *C* is below a given threshold *T. T* is initially set to a small number (10). We then refine the initial sequence clusters through the following steps: (1) A consensus sequence for each cluster is generated using partial order alignment^17^ (POA) on read sequences within each cluster. The consensus is used to update the cluster centroid. (2) After updating the centroids for all clusters, we again sort the clusters in order of the length of their revised centroids. (3) We iterate through the clusters, merging two clusters whenever the edit distance between their updated centroids is below *T*. (4) This process is repeated until no further clusters can be merged. (5) After cluster refinement, we only include clusters with a number of reads above *min*(10, 0.1 × n_s_) where *n*_*s*_ is the number of excluded reads per sample *s*. Finally, we check if the total number of reads in all remaining clusters are more than 0.8 × n_s_, otherwise we will increase *T* (attempting in order the following values: 20, 50, 80, 100, 150, 200, 300, 400, 500, 600, 700) to relax the constraints on sequence similarity for both adding sequences to existing clusters and merging cluster and repeat the steps above. Centroid sequences of final clusters are added to the set of potential haplotypes. Edit distance is calculated using Needleman-Wunsch algorithm with parameters gap_score = 1, match_score = 0, mismatch_score = 1. We optimized the algorithm by computing a lower bound of total edit distance at each row of the dynamic programming table *dp*[*n*][*m*], where *n* and *m* are length of sequences. For each cell in a row *i*, we iterate over columns *j* and compute min(dp[*i*, j] + abs((n − m) − (*i* − j))). This value shows the minimum number of insertions or deletions needed to reach cell *dp*[*n, m*]. If the minimum value per row exceeds the threshold *T*, computation stops.

#### Alignment of reads to candidate haplotypes

The original HipSTR approach employs a rigid alignment technique for aligning repeat sequences to candidate haplotypes, operating under the assumption that errors occur in multiples of the repeat unit length and happen only once within the repeat. However, these assumptions do not hold true for long reads (**Supplementary Fig. 1**), likely due to the fact that errors are driven by other processes than PCR. Instead, numerous errors can be present at different positions of the repeat rather than being strictly tied to the repeat unit size (**Supplementary Fig. 2**), with single base insertions or deletions most prevalent. To allow for a more flexible error model, we used a Hidden Markov Model approach based on that of Dindel^18^ to align reads to each candidate haplotype.

LongTR models the error in homopolymer repeats using a geometric distribution as follows:

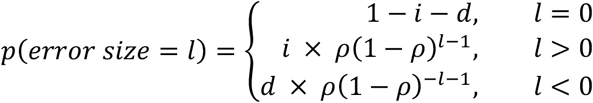

Where *ρ* controls the size of error, *i* and d are the probability of error increasing or decreasing the length of repeat respectively. To obtain the values for *i, d*, and *ρ*, we used the ALLREADS format field in LongTR output to extract the observed read lengths at each locus and use assembly genotypes as ground truth to compute the base pair differences *s* with actual genotype. Then 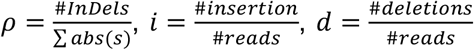were computed for homopolymers falling in each specific length range.

#### Phasing information

We leverage HipSTR’s existing option to use read phase information from haplotagged reads (and renamed the option from --10x-bams to --phased-bams) to accurately identify sample haplotypes. To consider phasing information, LongTR requires that at least one read from each haplotype be present, and unphased reads constitute no more than 20% of the total sample’s reads.

#### Genotyping

LongTR iterates over all pairs of haplotypes (*H*_*i*_, *H*_j_) (including homozygous pairs) and calculates a score for each possible genotype *G*(*H*_*i*_, *H*_j_) based on all observed reads *R* using the following formula:

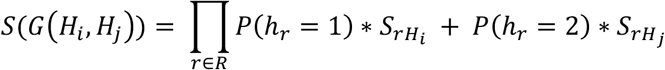

Where 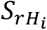 is the alignment score from aligning read *r* to haplotype *H*_*i*_ and *p*(*h*_*r*_ = 1) is the probability that read *r* is generated from the first haplotype. When read phase information is available, *p*(*h*_*r*_ = *k*) is set to 0 or 1 based on the haplotag field. Otherwise we set the probability to come from haplotype 1 vs. 2 as equally likely. The score for each haplotype pair is then normalized by the sum of scores for all possible genotypes and the result is reported as the quality score of the genotype in LongTR output, defined as Q.

#### Implementation

LongTR is implemented in C++. It leverages the HTSlib^19^ library to read directly from cloud addresses or URLs, which can avoid the costly step of downloading large sequence alignment files.

### Benchmarking against TRGT

TRGT v0.4.0 was used to genotype TRs from PacBio HiFi reads using non-default parameters --karyotype XY, --flank-len 25, and --max-depth 10000. The TRGT reference set was downloaded from TRGT GitHub (see **Data Availability**) and was used to perform genotyping with both TRGT and LongTR. To evaluate repeat allele sequences returned by each method, the HG002 assembly v1.0.1^12^ (**Data Availability**), with higher accuracy at homopolymers, was mapped to the GRCh38 reference genome using minimap2^20^ v2.26-r1175. For each repeat on autosomal chromosomes, allele sequences from the maternal and paternal haplotypes that completely span the repeat were extracted and compared to allele sequences reported in the output VCF files of LongTR and TRGT. For Mendelian Consistency analysis on autosomal chromosomes, LongTR was used to perform joint genotyping in all three samples and TRGT was run separately on each sample. We only considered a repeat in the Mendelian consistency analysis if 1) all samples were successfully genotyped at that repeat and 2) the genotype for at least one sample was not homozygous for the reference allele. The minimum quality score reported by LongTR among all three samples is considered the assigned score for that repeat. MI analysis was performed on autosomal chromosomes.

### Evaluation on ONT Duplex reads

Oxford Nanopore Duplex data for HG002 (**Data Availability**) was aligned to GRCh38 using minimap2 v2.26-r1175. LongTR v0.0.1 was then run on the TRGT reference restricting to autosomal chromosomes.

### Benchmarking against adVNTR

For evaluation of adVNTR and LongTR on VNTRs, adVNTR v1.5.0 was run with non-default parameters -- accuracy-filter, --pacbio, and --log-pacbio-reads. LongTR was run with non-default parameters --min-reads 4, -- max-tr-len 10000, --indel-flank-len 25, --phased-bam. The reference set of VNTRs was downloaded from the adVNTR GitHub (see **Data Availability**). Since adVNTR represents alternative alleles as integer multiples of the consensus repeat unit, direct allele length comparison was not possible. Therefore, we computed the concordance between LongTR and adVNTR copy number estimates allowing for one copy number difference to accommodate the complex VNTRs consisting of multiple motifs with different sequences. This analysis was done on autosomal chromosomes.

### Comparison to short read STR calls

HipSTR v0.7 was used to genotype STRs from 250bp paired-end PCR-free Illumina reads for HG002 (**Data Availability**) with non-default parameters --min-reads 4, --def-stutter-model, and --haploid-chrs chrY,chrX to genotype repeats with at least 4 overlapping reads, use default values for stutter error model, and to consider chromosome Y and X as haploid chromosomes. We ran HipSTR on PacBio HiFi reads with additional non-default parameters --use-unpaired, --no-rmdup, --max-str-len 10000, and --max-flank-indel 1 to use unpaired reads, not remove PCR duplicates, consider repeats with length up to 10000bp and allow for repeats with InDels found in their flanking regions.

## Code Availability

LongTR is open source and can be found on GitHub: https://github.com/gymrek-lab/LongTR.

## Data Availability

TRGT reference set of TRs: https://zenodo.org/record/8329210/files/adotto_repeats.hg38.bed.gz?download=1.

PacBio HiFi reads aligned to GRCh38 for HG002, HG003, and HG004: https://downloads.pacbcloud.com/public/revio/2022Q4/

PacBio HiFi reads for patients with pathogenic expansions and metadata: https://downloads.pacbcloud.com/public/dataset/RepeatExpansionDisorders_NoAmp.

HG002 assembly:

https://s3-us-west-2.amazonaws.com/human-pangenomics/T2T/HG002/assemblies/hg002v1.0.1.fasta.gz Oxford Nanopore Duplex data for HG002:

https://human-pangenomics.s3.amazonaws.com/index.html?prefix=submissions/0CB931D5-AE0C-4187-8BD8-B3A9C9BFDADE--UCSC_HG002_R1041_Duplex_Dorado/Dorado_v0.1.1/stereo_duplex/

adVNTR reference set of repeats: https://drive.google.com/file/d/1DetpBQySPNe2YAJa4FsjHn9qiRNS3wEV/view

Illumina reads for HG002:

https://ftp-trace.ncbi.nlm.nih.gov/ReferenceSamples/giab/data/AshkenazimTrio/HG002_NA24385_son/NIST_Illumina_2x250bps/novoalign_bams/.

HipSTR hg38 reference TR set:

https://github.com/HipSTR-Tool/HipSTR-references/raw/master/human/hg38.hipstr_reference.bed.gz

