## Supplementary Text for "Genome-wide profiling of genetic variation at tandem repeat from long reads"

### Supplementary Figures

#### Supplementary Figure 1

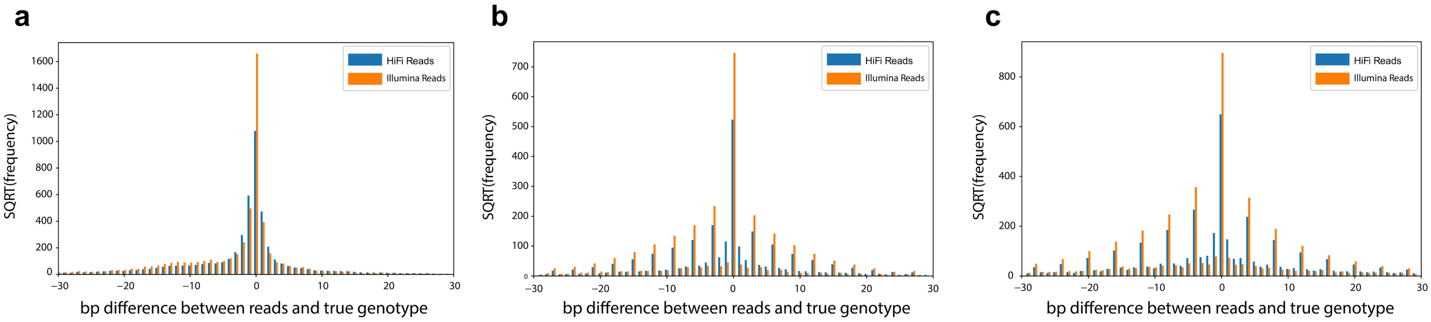


**Base pair error distribution for PacBio HiFi and Illumina reads.** Bars show the square root of the frequency of error sizes for PacBio HiFi (blue) and Illumina (orange) reads from sample HG002. **(a)** bp difference with the maximum likelihood genotype for homopolymers with total length between 20-60bp; **(b)** bp difference with the maximum likelihood genotype for trinucleotides with total length between 30-100bp; **(c)** bp difference with the maximum likelihood genotype for tetranucleotides with total length between 40-100bp. Notably, for **b-c**, PacBio reads show a higher rate of errors that are not a multiple of the repeat unit length.

**Supplementary Figure 2**

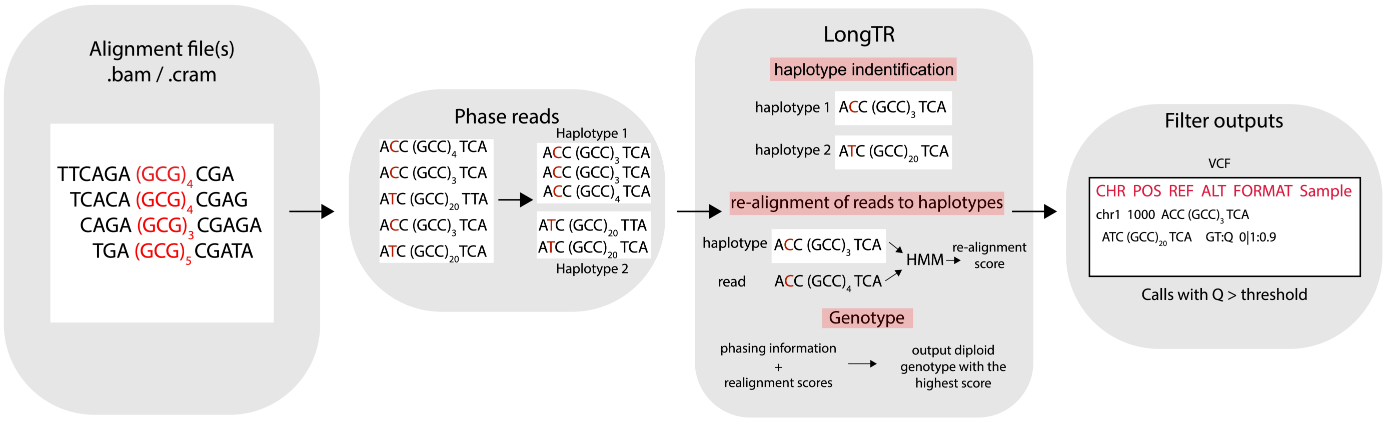


**LongTR workflow**. Users may first optionally haplotag (phase) the input reads. The aligned reads, along with a reference set of TRs, are input to LongTR for genotyping. LongTR uses a clustering strategy combined with partial order alignment to infer consensus haplotypes from error-prone reads, followed by sequence realignment using a Hidden Markov Model to infer the highest scoring genotypes. Finally, it outputs a VCF file with inferred genotypes, quality scores, and other fields that can be used to further filter low quality calls.

**Supplementary Figure 3**

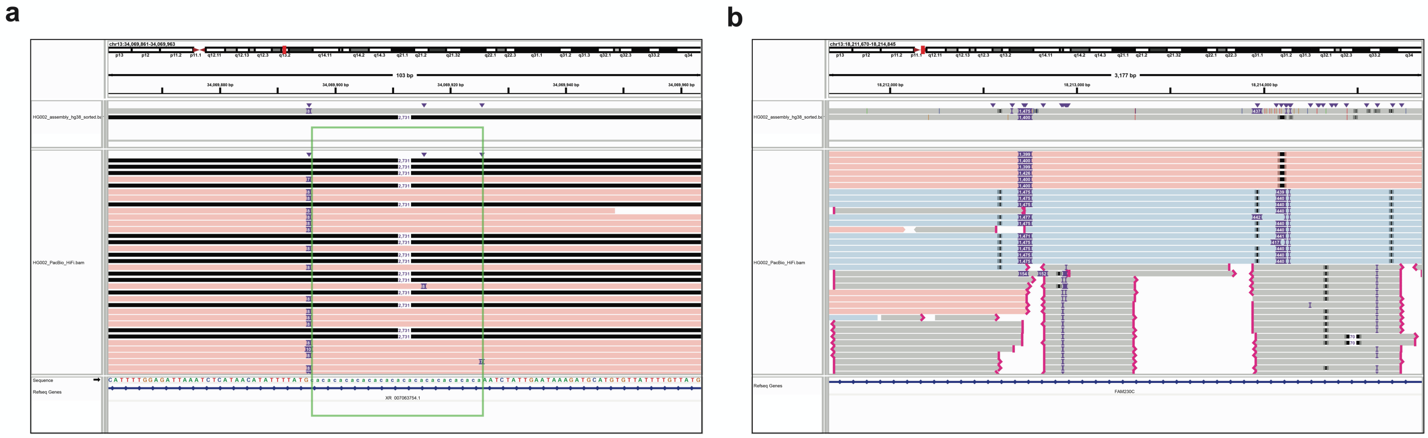

**IGV screenshots of examples where LongTR matches the assembly but TRGT does not.** The top window shows alignments of the maternal and paternal assemblies and the bottom window shows aligned HiFi reads to GRCh38. Red and blue denote PacBio HiFi reads from the two haplotypes of HG002 based on haplotag information. Gray reads have no haplotag information. The screenshot was generated using the Integrative Genomics Viewer[1]. **(a)** LongTR detects structural deletions resulting in null TR alleles**.** The repeat boundary is denoted by the green box. Note no reads from the haplotype labeled in blue are visible since those reads harbor a 2,731bp deletion at this locus (denoted by the black lines). **(b)** The window shows the repeat boundary. TRGT reports deletion in regions with high number of truncated reads.

**Supplementary Figure 4**

**a** **b**

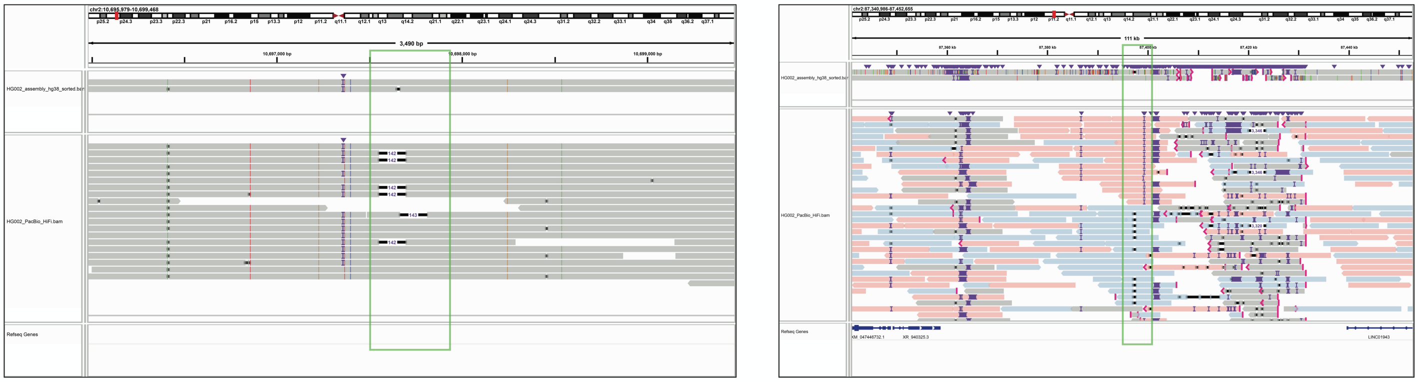

**IGV screenshots comparing HG002 assembly alleles with PacBio HiFi reads aligned to GRCh38 at loci where TRGT and LongTR agree but do not match the assembly.** The top window shows alignment of the maternal and paternal assemblies and the bottom window shows aligned HiFi reads. Red and blue denote PacBio HiFi reads from the two haplotypes of HG002 based on haplotag information. Gray reads have no haplotag information. The repeat boundary is denoted by the green box. **(a)** shows a VNTR with a 141bp repeat unit and total length of 474bp on GRCh38. Example of a highly homozygous region where assembly alleles are most likely incorrect as neither contain the 142bp deletion supported by multiple reads. **(b)** shows a VNTR with repeat unit of CCAAGCCAG and total length of 5,180bp on GRCh38 where the insertion on the red haplotype is missing from the assembly due to a nearby highly divergent tandem repeat that causes a break in assembly alignment.

**Supplementary Figure 5**

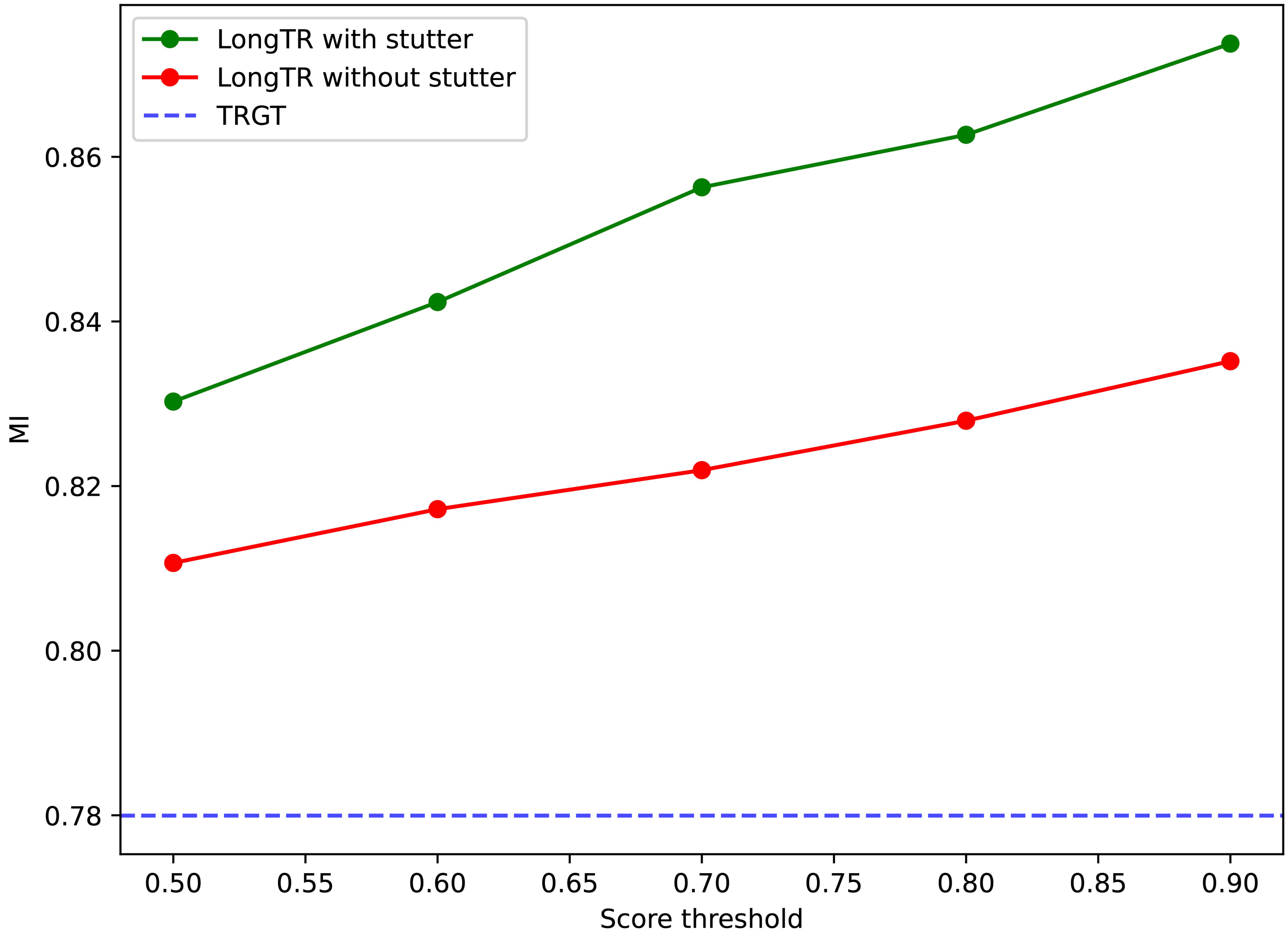


**Mendelian Inheritance for an Ashkenazi Trio at homopolymer TRs as a function of LongTR quality score.** The x-axis shows LongTR quality score threshold, and the y-axis shows the percent of genotyped loci that follow Mendelian Inheritance. The green line shows LongTR with stutter error modeling, the red line shows LongTR without using stutter error modeling, and the blue dashed line shows TRGT. Each locus was included if all 3 samples passed the score threshold. Loci where all three samples were homozygous for the reference allele were excluded from the analysis.

#### Supplementary Figure 6

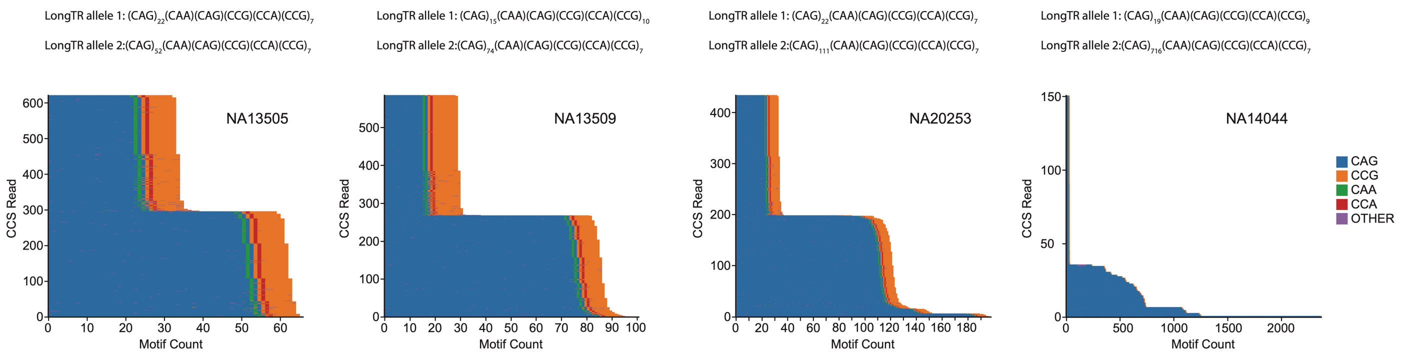


**Waterfall plots for reads from samples with an *HTT* expansion genotyped by LongTR**. Each row in each plot represents a single HiFi read. The x-axis shows the count of each motif identified in each read. LongTR precisely identified heterozygous allele sequences for which the repeat unit with the expansion can be inferred. From the left, plots show reads for samples NA13505, NA13509, NA20253, and NA14044. Plots were generated using TRviz[2].

**Supplementary Figure 7**

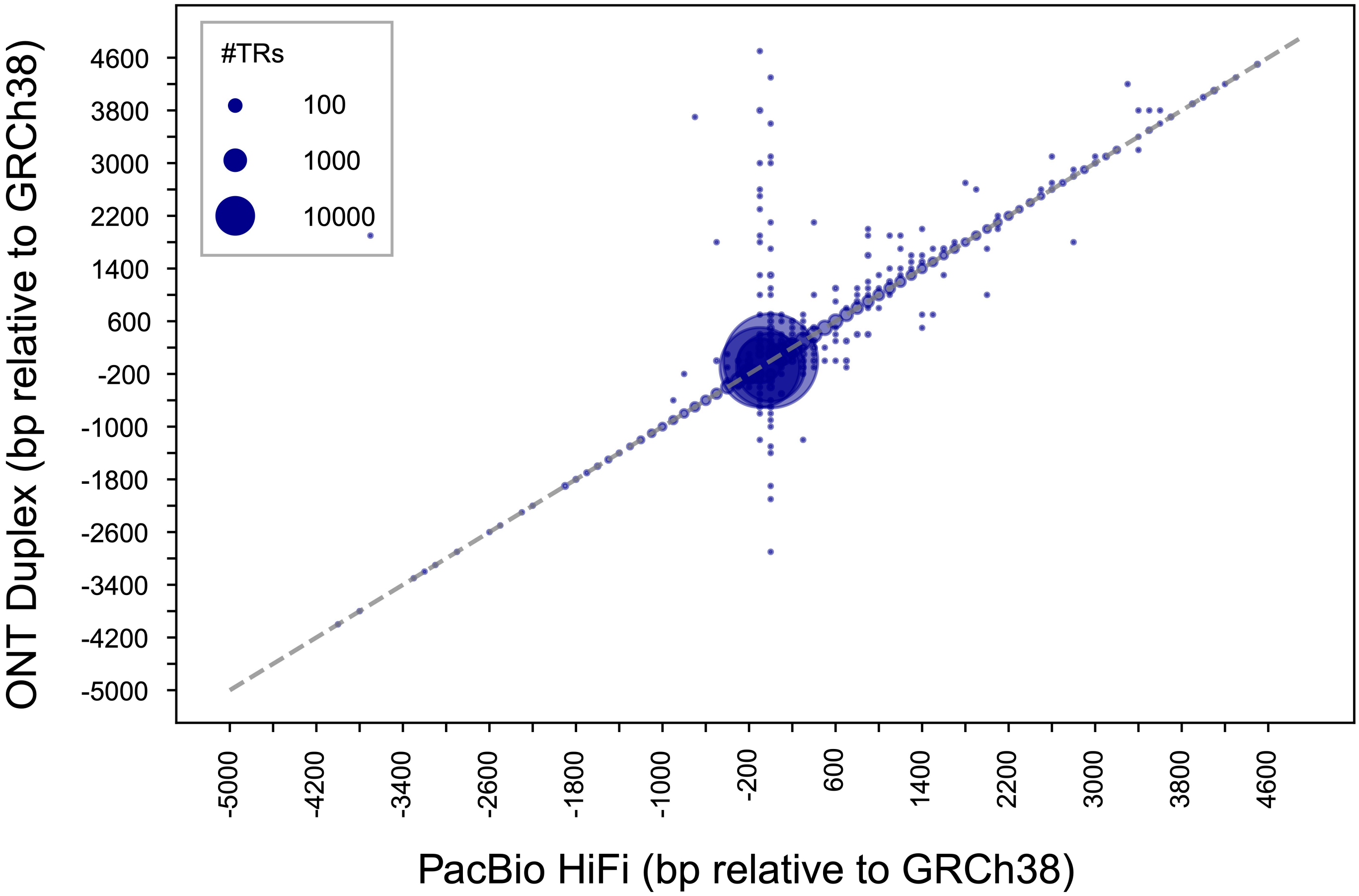


**Comparison of LongTR genotypes on ONT Duplex vs. PacBio HiFi data.** For each call, we computed the average of the length of each allele relative to the GRCh38 reference. The x-axis gives the calls using PacBio data and the y-axis gives the calls using ONT Duplex data. Bubble size scales with the number of calls at each coordinate.

#### Supplementary Figure 8

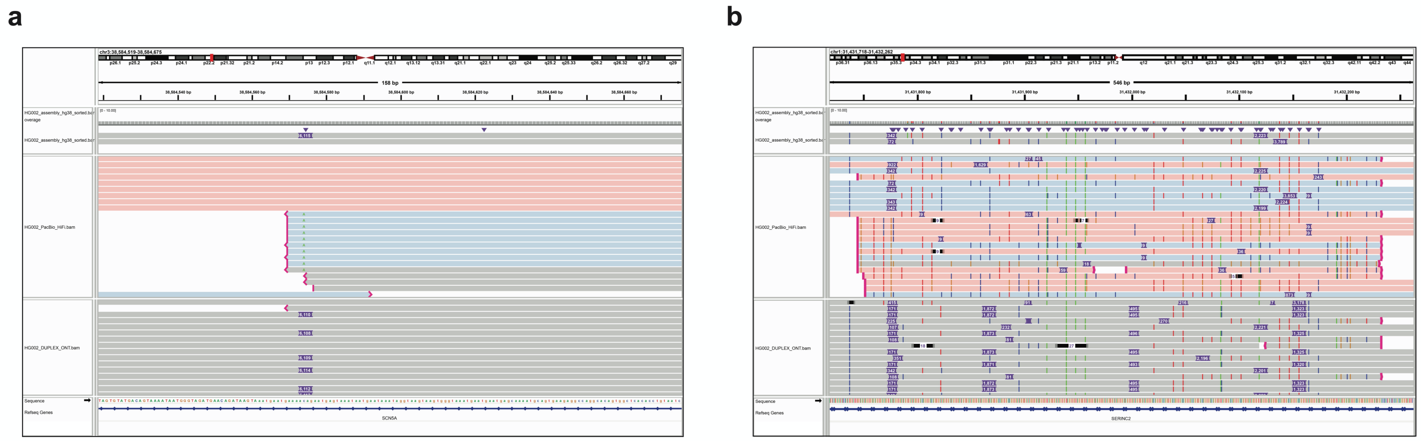

**IGV screenshots comparing LongTR calls on PacBio HiFi reads vs. ONT duplex reads.** The top window shows the assembly alignment, the middle window shows aligned HiFi reads, and the bottom window shows aligned ONT duplex reads. All data is aligned to GRCh38. Windows show the repeat boundary. Red and blue denote PacBio HiFi reads from the two haplotypes of HG002 based on haplotag information. Gray reads have no haplotag information. **(a)** shows a (AGTAAATAATG)n VNTR of total length of 156bp. Example of an insertion where aligned HiFi reads were clipped, resulting in an incorrect genotype call. **(b)** shows a (GGTGGATAG)n VNTR with total length of 544bp. Example of a heterozygous insertion where phased PacBio HiFi reads from the red haplotype are missing the insertion. In both cases when using ONT duplex reads LongTR was able to identify the insertion alleles.

**Supplementary Figure 9**

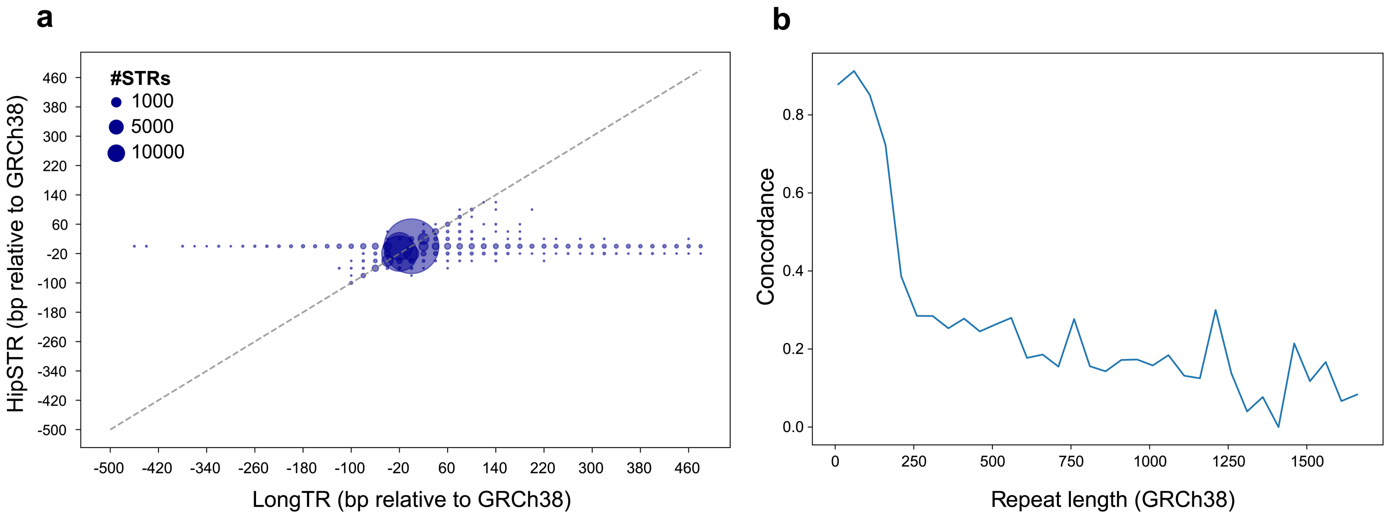


**Comparison between STR calls by HipSTR using Illumina short reads vs. LongTR on PacBio HiFi reads from sample HG002. (a) Length concordance between HipSTR (short reads) vs. LongTR (long reads) genotypes**. The x-axis and y-axis show the average base pair difference from the GRCh38 reference genome for the two alleles at each TR. Bubble size shows the number of points at each coordinate. **(b) Most discordances are at longer TRs.** The x-axis shows the repeat length on hg38. The y-axis shows the fraction of concordant calls between HipSTR and LongTR.

### Supplementary Tables

#### Supplementary Table 1

**Inferred error model parameters at homopolymer TRs in PacBio HiFi reads from HG002.** Parameters u and d indicate the probability to see an expansion or deletion error in each read. The step size of errors (length difference from the true allele in bp) is characterized by a geometric distribution with a parameter 'p'. We computed separate error models for different length ranges of homopolymers. As expected, errors become larger and more frequent with the increase in homopolymer repeat length.

| Homopolymer repeat length (bp) | Step-size (p) | Deletion coefficient (d) | Insertion coefficient (u) |
|---|---|---|---|
| 10-19 | 0.92 | 0.15 | 0.10 |
| 20-29 | 0.86 | 0.23 | 0.13 |
| 30-39 | 0.80 | 0.28 | 0.14 |
| 40-49 | 0.75 | 0.31 | 0.14 |
| >=50 | 0.71 | 0.34 | 0.15 |

**Supplementary Table 2**

Base pair differences from the reference genome at four TR loci were measured by LongTR using PacBio HiFi data. The first four patients have expansions in *HTT*. The next three have expansions in *FMR1*. HEK293 has no known expansion at these loci. None of the samples have expansions in *ATXN10* or *C9orf72*.

| Sample | HTT | | ATXN10 | | C9orf72 | | FMR1 | |
|---|---|---|---|---|---|---|---|---|
| | TRGT | LongTR | TRGT | LongTR | TRGT | LongTR | TRGT | LongTR |
| **NA12505** | 9,96 | 9,96 | 0,20 | 0,20 | 24,42 | 24,42 | 30,33 | 30,33 |
| **NA13509** | -3,168 | -3,168 | -10,30 | -10,30 | 12,18 | 12,18 | 30,33 | 30,33 |
| **NA20253** | 9,273 | 9,276 | -5,10 | -5,10 | -6,-6 | -6,-6 | 0,0 | 0,0 |
| **NA14044** | 6,1876 | 6,2115 | -10,-10 | -10,-10 | -6,-6 | -6,-6 | 30,33 | 30,33 |
| **NA13664** | -6,0 | -6,0 | 0,0 | 0,0 | -6,-6 | -6,-6 | 30,99 | 30,99 |
| **NA06896** | -18,3 | -18,3 | -5,20 | -5,20 | -6,30 | -6,30 | 9,509 | 9,426 |
| **NA07537** | -18,-6 | -18,-6 | -10,10 | -10,10 | -6,-6 | -6,-6 | 27,954 | 27,1000 |
| **HEK293** | -6,-3 | -6,-3 | 10,10 | 10,10 | -6,6 | -6,6 | 30,33 | 30,30 |
